# Aiptasia as a model to study metabolic diversity and specificity in cnidarian-dinoflagellate symbioses

**DOI:** 10.1101/223933

**Authors:** Nils Rädecker, Jean-Baptiste Raina, Mathieu Pernice, Gabriela Perna, Paul Guagliardo, Matt R Kilburn, Manuel Aranda, Christian R Voolstra

## Abstract

The symbiosis between cnidarian hosts and microalgae of the genus *Symbiodinium* provides the foundation of coral reefs in oligotrophic waters. Understanding the nutrient-exchange between these partners is key to identifying the fundamental mechanisms behind this symbiosis. However, deciphering the individual role of host and algal partners in the uptake and cycling of nutrients has proven difficult, given the endosymbiotic nature of this relationship. In this study, we highlight the advantages of the emerging model system Aiptasia to investigate the metabolic diversity and specificity of cnidarian – dinoflagellate symbiosis. For this, we combined traditional measurements with nano-scale secondary ion mass spectrometry (NanoSIMS) and stable isotope labeling to investigate carbon and nitrogen cycling both at the organismal scale and the cellular scale. Our results suggest that the individual nutrient assimilation by hosts and symbionts depends on the identity of their respective symbiotic partner. Further, *δ*^13^C enrichment patterns revealed that alterations in carbon fixation rates only affected carbon assimilation in the cnidarian host but not the algal symbiont, suggesting a ‘selfish’ character of this symbiotic association. Based on our findings, we identify new venues for future research regarding the role and regulation of nutrient exchange in the cnidarian - dinoflagellate symbiosis. In this context, the model system approach outlined in this study constitutes a powerful tool set to address these questions.

## Introduction

Coral reefs thrive in nutrient poor waters (1) and their ecological success fully relies on the nutrient-exchange between cnidarians and dinoflagellate algae of the genus *Symbiodinium* living in the host’s tissues (2, 3). In this association, the endosymbiotic algae translocate the majority of their photosynthetically-fixed carbon to the host, which in turn provides inorganic nutrients from its metabolism to sustain algal productivity (2, 4–6). The efficient recycling of organic as well as inorganic nutrients within this symbiosis underpins the high productivity of coral reefs in the absence of major sources of allochthonous nutrients (7, 8). Yet, this ecosystem is in global decline as anthropogenic environmental change impedes the role of cnidarians as key ecosystem engineers (9). Mass bleaching events, i.e. the disruption of cnidarian - dinoflagellate symbiosis signified by the expulsion of symbionts and physical whitening of corals on broad scales, are among the dominant drivers of this decline (10, 11). Understanding the causes of this symbiotic breakdown requires considering these symbiotic organisms as holobionts. Holobionts constitute complex metaorganisms that arise from the interaction of the hosts and their associated microorganisms such as protists, bacteria, and archaea (12). A crucial attribute of cnidarian holobionts is the ability to take up, assimilate, and exchange nutrients (13). In particular, nitrogen cycling appears to be key to the functioning of these holobionts (14, 15), since growth of *Symbiodinium* is nitrogen-limited in a stable symbiosis (14, 16–18). Nitrogen limitation might stabilize symbiont populations and facilitate the translocation of photosynthates to the host (19), a process providing most of the energy required for the host’s metabolism (2, 20). Yet, it is unclear whether the host can exert control over this translocation of nutrients (21–23).

Despite the importance of the individual contribution of host and symbionts to holobiont nutrient cycling (14, 23–25), studying these processes in scleractinian corals has proven difficult due to the complex and interwoven nature of the coral holobiont. As most corals are associated with a diverse *Symbiodinium* community and cannot be maintained in a symbiont-free stage, identifying underlying processes within these symbiotic interactions is challenging. In contrast, the emerging model organism Aiptasia (*sensu Exaiptasia pallida* (26)) offers distinct advantages to study the cnidarian-dinoflagellate symbiosis (27–29): (I.) this sea anemone can be reared in clonal lines, enabling the study of processes in the absence of biological variation (30); (II.) animals can be maintained in a symbiont-free stage, allowing to study host processes in the absence of symbionts (29); (III.) symbiont-free Aiptasia can be re-infected with specific symbiont strains, enabling the comparison of different symbionts (including those commonly associated with corals) in the same host background *in hospite* (31); (IV.) an extensive array of genetic resources is available in Aiptasia, allowing to link genetic and physiological traits (27). These distinct advantages will prove especially powerful to study metabolic interactions between host and symbionts, particularly if combined with state of the art imaging techniques such as nano-scale secondary ion mass spectrometry (NanoSIMS), which allow precise quantification of element distribution at high spatial resolution (32, 33). Coupled with stable isotope labeling, this technology enables imaging of metabolic processes at subcellular resolution and consequently quantification of nutrients assimilation at the single-cell level for each symbiotic partner (33). NanoSIMS has opened doors to an unprecedented level of information across all fields of biology and has been successfully applied to corals (34–38). In this study, we combined the advantages of the Aiptasia model system with high resolution NanoSIMS to showcase the advantages of this model approach for the study of metabolic interactions and diversity in the cnidarian – dinoflagellate symbiosis.

## Material & methods

### *Maintenance of* Aiptasia

Four different host–symbiont pairings were maintained in separate batches. These combinations involved two different host clonal lines (CC7 (41) and H2 (39)) as well as two different symbiont populations (A4 and B1 dominated (68)). Whilst CC7 Aiptasia can form stable associations with a diversity of *Symbiodinium* types, H2 Aiptasia show high fidelity to their native symbionts suggesting a higher selectivity and/or specificity with their symbionts (40). This specificity of H2 Aiptasia hinders reinfection with other symbionts thereby preventing a full factorial design in this study. Nevertheless, these host clonal lines provide an ideal basis for the comparison of symbiont diversity and specificity.

To allow comparison of symbiont types within the same host line and to compare performance of the same symbiont type within different host lines, CC7 Aiptasia were bleached and reinfected with type B1 (strain SSBO1) symbionts, previously isolated from H2 Aiptasia. For this, non-symbiotic CC7 Aiptasia were generated and reinfected as described by Baumgarten *et al*. (27). In brief, animals were repeatedly bleached by incubation in 4°C sterile seawater for 4 h, followed by 1–2 days at 25°C in sterile seawater containing the photosynthesis inhibitor diuron. Non-symbiotic animals were maintained for at least 1 month prior to reinfection to confirm absence of residual symbionts. For reinfection, non-symbiotic animals were subjected to three cycles of incubation for one day in sterile seawater containing 10^5^ *Symbiodinium* cells mL^−1^ followed by *Artemia salina* nauplii feeding the next day. Thus, the four combinations were: non-symbiotic CC7 Aiptasia, CC7 Aiptasia with its native A4 symbionts; CC7 Aiptasia reinfected with B1 symbionts and H2 Aiptasia with its native B1 symbionts (Fig. 1A-D). Animals were reared in sterile seawater (35 PSU, 25 °C ,~80 μmol photons m^−2^ s^−1^ on a 12h:12h light:dark schedule) and fed with freshly hatched *Artemia salina* nauplii three times per week. Animal cultures were propagated under these conditions for more than one year to ensure anemones recovered from bleaching and reinfection procedures and to confirm the stability of native and introduced symbiotic associations. Stability of *Symbiodinium* communities was monitored using qPCR as outlined by Correa *et al*. (69). Any feeding was abandoned three days prior to measurements to exclude potential confounding effects. Thereby this experimental design allowed us to disentangle the contribution of host and symbionts to holobiont nutrient cycling in three interesting comparisons: (1.) between different symbionts within the same host line, (2.) between different hosts lines with the same symbiont, and (3.) between symbiotic and non-symbiotic states within the same host line.

**Fig. 1.**
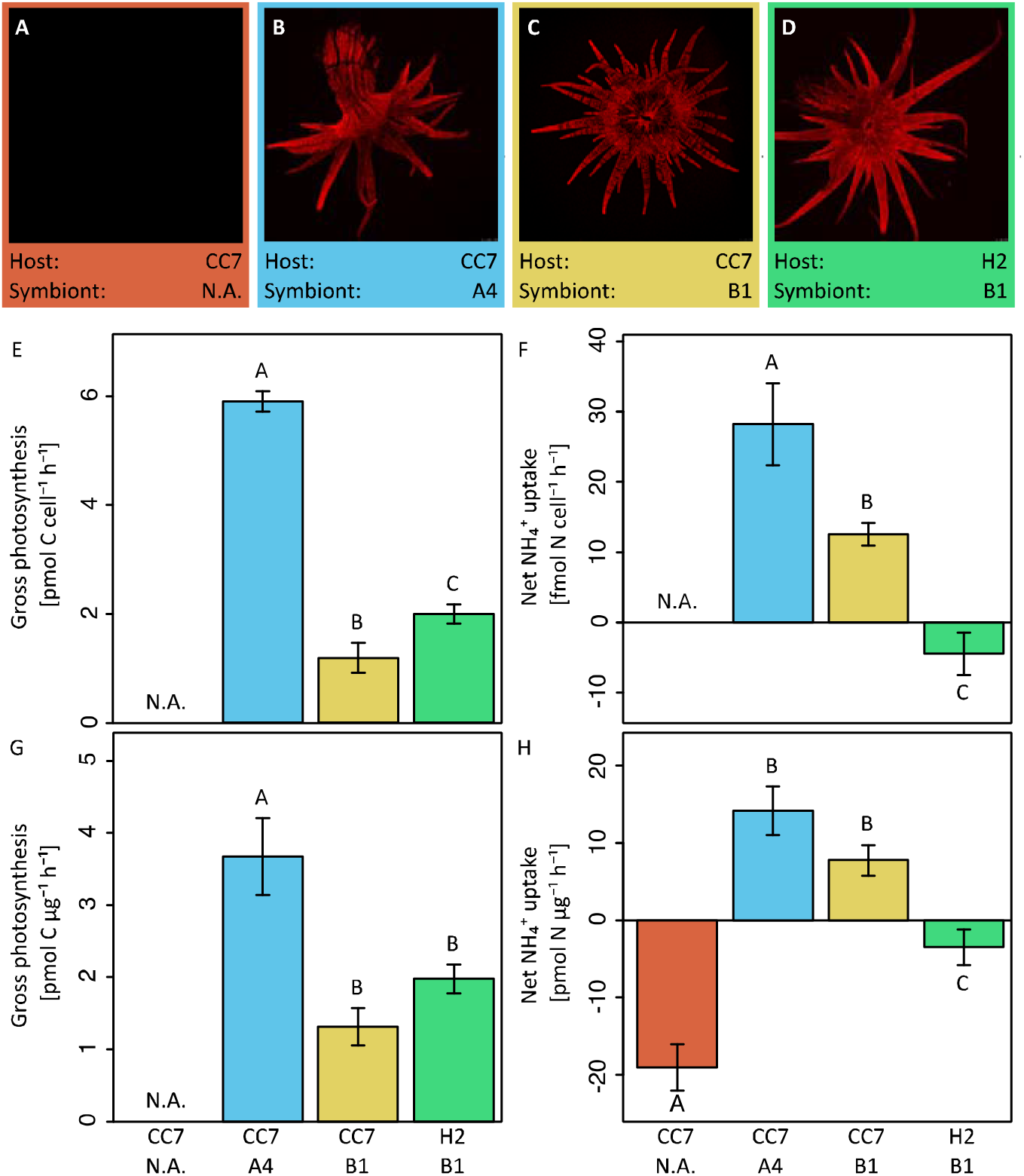
Ammonium (NH_4_^+^) uptake and carbon fixation (gross photosynthesis) of Aiptasia. Fluorescence microscopy overview of the four host–symbiont combinations **(A-D)** to visualizes chlorophyll autofluorescence of *endosymbiotic Symbiodinium*. Gross photosynthesis **(E,G)** and net NH_4_^+^ uptake **(F,H)** of Aiptasia were normalized either to symbiont density **(E,F)** or total host protein content **(G,H).** Gross photosynthesis rates were calculated as the sum of net photosynthesis and respiration rates (P_G_ =P_N_ + R). Net NH_4_^+^ uptake was quantified with the ammonium depletion method. All data shown as mean ± SE. Different letters above bars indicate significant differences between groups (p < 0.05).

### Oxygen flux measurements

Net photosynthesis and respiration rates were measured via oxygen (O_2_) evolution and consumption measurements during light and dark incubations, respectively. For this purpose, five specimens of each host–symbiont combination were transferred into 25 ml glass chambers filled with sterile seawater. Specimens were left to settle for 30 min in the dark, before magnetic stirrers were turned on to prevent stratification of the water column. Subsequently, O_2_ concentrations were recorded once per second over the course of 30 min incubations during the light (~80 μmol photons m^−2^ s^−1^, 25°C) and dark (<1 μmol photons m^−2^ s^−1^, 25°C) using FireSting O2 optical oxygen meters (PyroScience, Gemany). Following incubation all specimens were immediately flash frozen and stored at −20°C until further analysis. Net photosynthesis (inferred from light incubations) as well as respiration (inferred from dark incubations) rates were corrected for seawater controls and normalized to total protein content and *Symbiodinium* densities of specimens. O_2_ fluxes of net photosynthesis and respiration rates were transformed into their carbon equivalents using the photosynthetic and respiration quotients of 1.1. and 0.9 as proposed by Muscatine *et al*. (4). Gross photosynthesis rates were calculated according to: gross photosynthesis = net photosynthesis + |respiration| rate.

### Quantification of NH_4_^+^ uptake and release

Net uptake rates were assessed on the holobiont levels during light (~80 μmol photons m^−2^ s^−1^, 25°C) and dark (<1 μmol photons m^−2^ s^−1^, 25°C) conditions using the depletion technique (70). Five specimens of each host–symbiont combination were incubated for 60 min in 25 ml chambers filled with NH_4_^+^-enriched artificial seawater (ASW) with a final concentration of 5 μM (71). 10 ml water samples were collected before and after the incubation, filtered (45 μm) and immediately analyzed for ammonium concentrations using an autoanalyzer (SA3000/5000 Chemistry Unit, SKALAR, Netherlands). Differences in NH_4_^+^concentrations were corrected for seawater controls and normalized to incubation time, total host protein content and *Symbiodinium* densities of specimens to obtain net uptake rates during both light and dark incubations.

### Protein content, Symbiodinium density, and chlorophyll concentrations

Frozen specimens were defrosted in 500 μl sterile saline water and homogenized using a Micro DisTec Homogenizer 125 (Kinematica, Switzerland). Aliquots of the homogenate were immediately analyzed for total protein content as well as symbiont concentrations. For total host protein content, *Symbiodinium* cells were removed by brief centrifugation and the supernatant was analyzed with the Micro BCA Protein Assay Kit (Thermo Scientific, USA) using 150 μl of 15× diluted tissue slurry as per manufacturer instructions. Likewise, *Symbiodinium* density was quantified with fluorescence assisted cell sorting (BD LSRFortessa, BD Biosciences, USA) using 100 μl of strained tissue slurry.

### Isotope labeling and sample preparation

To verify nitrogen and carbon assimilation rates on the holobiont level, an isotopic labeling experiment was conducted for subsequent Nanoscale secondary ion mass spectrometry (NanoSIMS) analysis. Individual specimens of each host–symbiont combination were incubated for 24 h (12h:12 light dark cycle) in 25 ml incubation chambers containing ASW. For isotopic enrichment, freshly prepared ASW, essentially free from bicarbonate and ammonium, was supplemented with NaH^13^CO_3_ (isotopic abundance of 99%) as well as ^15^NH_4_Cl (isotopic abundance of 99%) at a final concentration of 2mM and 5 μM, respectively (adapted from Harrison *et al*. (71)). Following incubation, all specimens were immediately transferred to a fixative solution (2.5% glutaraldehyde, 1 M cacodylate) and stored at 4°C until further processing (within 14 days).

Individual tentacles were collected from each anemone under a stereomicroscope for further sample preparation adapted after Pernice *et al*. (34) and Kopp *et al*. (46). First, samples were post-fixed for 1h at RT in 1% OsO_4_ on Sörensen phosphate buffer (0.1 M). Samples were dehydrated in a serious of increasing ethanol concentrations (50%, 70%, 90%, 100%) followed by 100% acetone. Tissues were then gradually infiltrated with SPURR resin of increasing concentrations (25%, 50%, 75%, 100%). Subsequently, tissues were embedded in SPURR resin and cut into 100 nm sections using an Ultracut E microtome (Leica Microsystems, Germany) and mounted on finder grids for Transmission Electron Microscopy (ProsciTech, Australia).

### NanoSIMS imaging

Gold-coated sections were imaged with the NanoSIMS 50 ion probe at the Center for Microscopy, Characterisation and Analysis at the University of Western Australia. Samples surfaces were bombarded with a 16 keV primary Cs^+^ beam focused to a spot size of about 100 nm, with a current of approximately 2 pA. Secondary molecular ions ^12^C^12^C^−^, ^12^C^13^C^−^, ^12^C^14^N− and ^12^C^15^N^−^ were simultaneously collected in electron multipliers at a mass resolution (M/ΔM) of about 8,000, enough to resolve the ^12^C^13^C^−^ from the ^12^C_2_^1^H^−^ peak and the ^13^C^14^N^−^ and ^12^C^15^N^−^ peaks from one another. Charge compensation was not necessary. Five images of different areas within the gastrodermis of the tentacle (25 – 45 μm raster with 256 × 256 pixels) were recorded for all targeted secondary molecular ions by rastering the primary beam across the sample with a dwell-time of 10–20 ms per pixel. After drift correction, the ^13^C/^12^C or ^15^N/^14^N maps were expressed as a hue-saturation-intensity image (HSI), where the color scale represents the isotope ratio. Image processing was performed using the ImageJ plugin OpenMIMS (National Resource for Imaging Mass Spectrometry, https://github.com/BWHCNI/OpenMIMS/wiki).

Enrichment of the isotope labels was quantified for 20 ROIs (circles of 2-10 μm) per category (symbiont cells, gastrodermal host tissue and gastrodermal vesicles) for each host–symbiont combination, and expressed using *δ*^13^C and *δ*^15^N notation. Gastrodermal host tissue was quantified in the form for ROIs placed adjacent to symbiont cells as clear cell boundaries were not always identifiable.

Unlabeled Aiptasia served as unlabeled controls. *δ*^13^C and *δ*^15^N enrichment was quantified as follows:

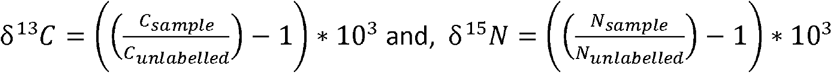

where N is the ^15^N/^14^N ratio of sample or unlabeled control and C is the ^13^C/^12^C ratio (measured as ^12^C^13^C^−^/^12^C^12^C^−^ Cions) of sample or unlabeled control, respectively. In this context, it is important to note that carbon and nitrogen incorporation at the cellular level was likely underestimated in our study as sample preparation for NanoSIMS may result in partial extraction of biomolecules.

### Statistical analysis

All statistical analyses were conducted with R version 3.2.2 (72). Data were tested for normal distribution using the Shapiro-Wilk test. All measurements on the holobiont level (gross photosynthesis, respiration, net NH_4_^+^ uptake) followed normal distribution and were analyzed with a one-way analysis of variance (ANOVA) using host-symbiont combination as explanatory variable; only gross photosynthesis rates normalized by symbiont density did not follow a normal distribution and hence were analyzed with a generalized linear model (GLM) using host-symbiont combination as the explanatory variable. Similarly, *δ*^13^C and *δ*^15^N enrichment data did not follow a normal distribution and were analyzed in two-factorial GLMs using additive as well as interactive effects of host-symbiont combination as well as holobiont compartment (host, lipid body, symbiont). All GLMs were fitted with Gamma distribution and ‘log’ function to optimize the fit of the model. Fit of model residuals were confirmed using the qqPlot() function as implemented in the ‘car’ package for R (73). An overview of model results is provided in the Supplementary Information Table S1. Adjustment for multiple comparisons between host-symbiont combinations and holobiont compartments was done following the Bonferroni procedure. Significant differences identified via the post hoc comparison are indicated in the figures as different letters above bars.

## Results

We investigated the relative contribution of cnidarian hosts genotypes and their dinoflagellate symbionts to assimilate dissolved inorganic nitrogen (as ammonium (NH_4_^+^)) and carbon (as bicarbonate) both at the organismal and at the cellular level in Aiptasia by assaying four different associations of hosts and symbionts (Fig. 1A-D). Taken together, these four host–symbiont combinations allowed us to identify nutrient dynamics in symbiotic and non-symbiotic Aiptasia and to address the following questions: (a.) whether different *Symbiodinium* types possess different metabolic capabilities within the same host strain and (b.) to what extent different host strains affect the metabolic performance of the same algal symbiont type.

### Carbon assimilation and translocation

Host–symbiont combination of Aiptasia showed distinct differences in carbon fixation both at the holobiont (Fig. 1E,G) as well as at the cellular level (Fig. 2A-H, see Supplementary Information Table S1 for an overview of statistical model results). While fixation rates were highly variable between the three groups of symbiotic Aiptasia, no carbon fixation was detectable in non-symbiotic Aiptasia, confirming that carbon assimilation was photosynthetically driven. At the holobiont level, gross carbon fixation (measured as gross photosynthesis) was highest in Aiptasia of the clonal line CC7 with their native symbiont community (*Symbiodinium* type A4) after normalization to symbiont density (Fig. 1E) or host protein content (Fig. 1G). In contrast, CC7 Aiptasia symbiotic with Clade B (SSBO1) *Symbiodinium* showed the lowest gross photosynthesis rates of all symbiotic Aiptasia groupings. In particular, rates were lower than H2 Aiptasia with the same type B1 dominated symbiont community. Photosynthetic carbon fixation was more than three-fold higher than dark respiratory carbon consumption in all symbiotic Aiptasia groupings. Overall dark respiration rates followed a less defined yet similar pattern as gross photosynthesis rates of host–symbiont combinations (Supplementary Information Fig. S1A,C), with animal holobionts showing a strong positive correlation between gross photosynthesis and respiration rates (Spearman's correlation, r_s_=0.930, p<0.001).

**Fig. 2.**
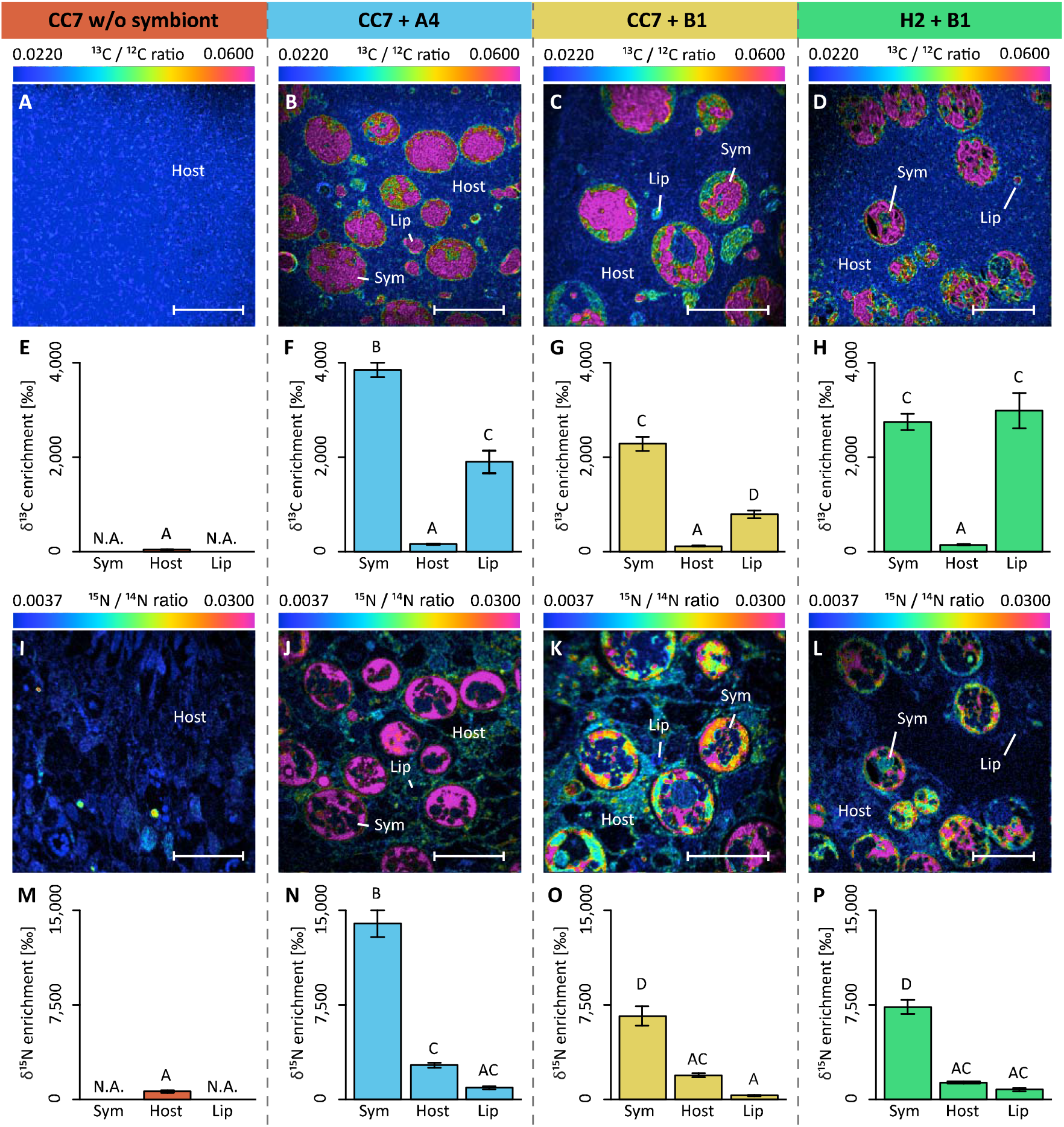
NanoSIMS imaging and quantification of cell specific carbon (as ^13^C-bicarbonate) and nitrogen (as ^15^N-ammonium) assimilation within the Aiptasia—*Symbiodinium* symbiosis. Representative images of the distribution of ^13^C/^12^C ratio **(A-D)** and of ^15^N/^14^N ratio **(l-L)** within the Aiptasia holobiont are displayed as Hue Saturation Intensity (HSI). The rainbow scale indicates the ^13^C/^12^C and ^15^N/^14^N ratio, respectively. Blue colors indicate natural abundance isotope ratios shifting towards pink with increasing ^13^C and ^15^N incorporation levels, respectively. Corresponding *δ*^13^C enrichment **(E-F)** and *δ*^15^N enrichment **(M-P)** in the tissues of the four host–symbiont combinations. For each NanoSIMS image, the *δ*^13^C (E-F) and *δ*^15^N (M-P) enrichment were quantified for individual Regions Of Interest (ROIs) that were defined in OpenMIMS by drawing (I) the contours of the symbionts, and circles covering (II) the adjacent host tissue and (III) the host lipid bodies. Scale bars represent 10 μm. Abbreviations: Sym = *Symbiodinium* cell, Host = tissue (host) & Lip = lipid body (host). All data shown as mean ± SE. Different letters above bars indicate significant differences between groups (p < 0.05).

Isotope labeling and NanoSIMS imaging revealed that these observed differences on the holobiont level translated into an intricate picture at the cellular level (Fig. 2A-H). First, *δ*^13^C enrichment was evident in both host and symbiont cells in all symbiotic Aiptasia groupings (Fig. 2B-D). Second, although enrichment was highest in *Symbiodinium* cells, localized regions of < 5 μm diameter in the host tissue (referred to as ‘lipid bodies’ from this point on) also showed significantly higher rates of enrichment compared to the surrounding host tissue. Third, Clade B *Symbiodinium* showed no differences in ^13^C-incorporation depending on the host, and incorporation rates were 30–40 % lower than in Clade A symbionts. Host lipid bodies, on the contrary, showed a reversed picture with Clade B associated H2 Aiptasia having the highest and Clade B associated CC7 Aiptasia having the lowest ^13^C assimilation rates, despite harboring the same symbiont types.

### NH_4_^+^ assimilation and release

Similar to carbon fixation, strong differences in ammonium (NH_4_^+^) assimilation were evident between the experimental groups of Aiptasia at both holobiont (***χ***_(3,16)_ = 87.44, p < 0.01) and cellular levels (***χ***(3,352) = 64.06, p < 0.01). At the holobiont level, all four host—symbiont combinations showed higher NH_4_^+^ uptake/release rates during the light (Fig. 1F,H), compared to dark conditions (Supplementary Information Fig. S1B,D). When normalized to host protein content, non-symbiotic Aiptasia showed the highest net release of NH_4_^+^ at the holobiont level both during light (Fig. 1H) and dark incubations (Supplementary Information Fig. S1D). Albeit significantly lower, symbiotic H2 Aiptasia also had a net release of NH_4_^+^ into the surrounding seawater during the light incubations. In contrast, both groups of symbiotic CC7 Aiptasia showed a net uptake of NH_4_^+^ by the holobiont during both light and dark conditions. Further, the uptake rate was affected by the associated symbiont community, with Clade A dominated CC7 holobionts taking up more NH_4_^+^ than their Clade B infected counterparts (Fig. 1F,H).

Although NH_4_^+^ assimilation ranged from net uptake to net release in the different experimental groups, NanoSIMS imaging confirmed that all four host–symbiont combinations incorporated ^15^N into their cells (Fig. 2I-P). Whilst *δ*^15^N signatures were highest in *Symbiodinium* cells, ^15^N assimilation was also observed within the cnidarian host tissue including that of non-symbiotic Aiptasia. Similar to *δ*^13^C patterns, *δ*^15^N enrichment in *Symbiodinium* cells aligned with algal symbiont type rather than host identity, and Clade B symbionts showed lower rates of incorporation than Clade A. Conversely, ^15^N incorporation into host cells was not significantly different between symbiotic Aiptasia groupings, irrespective of their symbiont type. Non-symbiotic CC7 Aiptasia had the lowest overall ^15^N incorporation into their tissue, yet showed small (< 5 μm in diameter) and localized regions of high enrichment. In contrast, the afore-mentioned lipid bodies of high *δ*^13^C-enrichment showed consistently lower *δ*^15^N-signatures than surrounding host tissues in all three symbiotic Aiptasia strains.

## Discussion

Aiptasia has proven to be a powerful emerging tool for the genetic and molecular study of the cnidarian – alga symbiosis (27, 41, 43). Beyond these realms, only few studies have begun to exploit the advantages that Aiptasia has to offer (24, 25, 44, 45). Here, we set out to assess the use of Aiptasia as a model to study nutrient cycling in the cnidarian – alga symbiosis. Whilst NanoSIMS has been successfully used previously to study nutrient uptake in corals (46–48), the flexibility of the Aiptasia model enables for the first time to decouple the relative contribution of the host and symbionts to nutrient cycling. Although the methodology outlined in our approach was optimized to trace carbon and nitrogen assimilation within coral or Aiptasia holobiont (34, 46), the method can be easily modified depending on the experimental requirements. Specific labeled compounds can also be used as tracers to follow the translocation and uptake of specific molecules in complex systems by coupling the spatial resolution of NanoSIMS with the molecular characterization afforded by time-of-flight secondary ion mass spectrometry (ToF-SIMS) (49). Also, as shown here, detailed cellular insights gained from NanoSIMS will prove most powerful when integrated with traditional holobiont based measurements to identify the complexity of processes. Using this integrative approach, differences in nutrient assimilation across different host–symbiont associations became evident, both at the holobiont as well as the cellular level. Yet, only the integration of both levels of biological organization allowed to comprehensively disentangle some of the intricacies of nutrient cycling in the Aiptasia holobiont.

### Carbon cycling in Aiptasia

All three groups of symbiotic Aiptasia showed high rates of gross photosynthesis that exceeded their respiratory carbon requirements thereby supporting net productivity of the holobiont required for stable symbiotic associations (4). Yet, differences in gross photosynthesis between host–symbiont combinations were evident at the holobiont level. Gross photosynthesis rates differed between the same host infected with different algal symbionts and between different hosts infected with the same algal symbionts. Thereby our findings support the findings by Stazark *et al*. (24) who reported differences in carbon flux depending on symbiont type and between heterologous and homologous symbionts in Aiptasia, confirming previous observations that carbon fixation depends on the interaction of both host and symbionts (24, 25, 36, 50).

At the cellular level, we observed particular areas of *δ*^13^C enrichment (hotspots) in the host tissue similar to previous observations (36, 51). This high *δ*^13^C enrichment is further coupled with lower *δ*^15^N enrichment, suggesting that these hotspots likely constitute a form of carbon storage compartments in the host tissue. Based on shape, size, and location in the tissue, these compartments are most likely lipid bodies (52). These cellular organelles are abundant in symbiotic cnidarians as they allow for rapid short-term carbon storage and remobilization depending on cellular carbon availability (53). Hence, amount, size and enrichment of these lipid bodies may be an excellent proxy to assess the amount of carbon translocated by *Symbiodinium* to the host, but further studies are needed to unequivocally determine their nature. Lipid body enrichment in the host was highest in H2 Aiptasia and lowest in CC7 Aiptasia, both associated with *Symbiodinium* type B1. Yet, *δ*^13^C enrichment in algal cells was unaffected by host identity. At the same time, our results revealed that Clade A and B symbionts had distinctly different *δ*^13^C enrichment, even in the same clonal Aiptasia host line. These differences are likely the consequence of differential metabolic requirements by the specific symbionts. Thus, *δ*^13^C enrichment may be a powerful tool to differentiate between symbiont types *in hospite*.

Taken together, observed differences in gross carbon fixation at the holobiont level were reflected in the combined *δ*^13^C enrichment (host tissue + lipid bodies + symbionts) at the cellular level. However, NanoSIMS data revealed that these patterns were only caused by differences in enrichment of the host lipid bodies (a proxy of carbon translocation to the host). In contrast, *δ*^13^C enrichment in algal cells differed depending on symbiont type (i.e., showed stable *δ*13C enrichment within the same symbiont types), but was unaffected by host identity. This apparent contradiction may have important implications for our understanding of symbiosis functioning. The fact that *δ*13C enrichment in algal cells differed only depending on symbiont type but was unaffected by host identity implies that symbionts retained the same amount of fixed carbon regardless of overall fixed carbon availability. Hence, only excess carbon, not consumed by algal metabolism, appears to be available for translocation to the host. Therefore, factors reducing the availability of excess carbon in the symbiont, may potentially deprive the host of its main energy source, despite harboring viable symbionts in its tissue. This ‘selfish’ aspect of the symbiosis may pose a potential threat to the stability of the holobiont under conditions of reduced fixed-carbon availability, such as those imposed by environmental stress (54, 55).

### Nitrogen cycling in Aiptasia

The observation of drastically different carbon fixation and translocation rates between different host–symbiont combinations raises questions regarding the underlying regulatory mechanisms of carbon cycling within these symbioses (13). Importantly, nitrogen availability *in hospite* has been proposed to be among the environmental controls of these processes (14, 15, 19, 56). Indeed, drastic differences in nitrogen assimilation became evident when comparing different host–symbiont combinations. Strikingly, the two different host lines Aiptasia H2 and CC7 showed net NH_4_^+^ release and NH_4_^+^ uptake during the light, respectively, even when hosting the same algal symbionts. These findings suggest that the *in hospite* nutrient availability for the symbiont may be drastically different depending on the associated Aiptasia host. Hence, differences in gross photosynthetic activity and translocation may be partly attributed to variations in availability of nitrogen derived from the host metabolism. Interestingly, while CC7 Aiptasia showed light-enhanced NH_4_^+^uptake as previously reported for corals (57), H2 Aiptasia showed net release of NH_4_^+^ during the light, contrasted by slight uptake during the dark. While we cannot explain this discrepancy at this point, it illustrates the drastic effects of host identity on nitrogen assimilation of the holobiont. At any rate, our results highlight the functional diversity and specificity of cnidarian-dinoflagellate symbioses, prompting research across a range of host–symbiont combinations.

At this point it is not possible to distinguish whether the increased *δ*^15^N enrichment of host tissues in symbiotic animals are due to direct NH_4_^+^fixation by the host or the translocation of fixed nitrogen by the symbiont. However, nitrogen assimilation was observed even in the absence of algal symbionts, as evidenced by non-symbiotic Aiptasia. Although these animals showed a high net release of NH_4_^+^ at the holobiont level, NanoSIMS imaging confirmed the incorporation of ^15^N within localized hotspots of their tissue at low rates. While the exact nature of these hotspots remains unknown at this point, our results confirm that Aiptasia also has the ability to assimilate inorganic nitrogen from seawater as previously reported for corals (34). However, it remains to be determined whether this capability is intricate to the host cellular machinery or a function of associated bacterial symbionts or both (48).

In contrast to *δ*^15^N enrichment of their hosts, *Symbiodinium* types showed characteristic *δ*^15^N enrichment patterns regardless of the identity of their host. Hence, *δ*^15^N enrichment may prove a useful tool to identify symbiont identity *in hospite*, especially when combined with *δ*^13^C measurements.

Different to carbon fixation measurement, patterns of NH_4_^+^ uptake on the holobiont level were not directly reflected in the overall *δ*^15^N enrichment on the cellular level. Specifically, symbiont-free CC7 Aiptasia as well as symbiotic H2 Aiptasia showed net release of NH_4_^+^ from the holobiont during light conditions, yet NanoSIMS analysis confirmed the incorporation of ^15^N from surrounding seawater. While these differences may be partly attributed to differences in incubation time and light availability for the two measurements, they further suggest that uptake and release of NH_4_^+^ appear to be in a dynamic equilibrium in Aiptasia. Hence, the stable *δ*^15^N enrichment of the same symbiont type in CC7 and H2 suggests that the contribution of nitrogen derived from host metabolism was negligible compared to the incorporation of nitrogen from seawater under these conditions. Under natural oligotrophic conditions, however, host metabolism may make a significant contribution to the nitrogen supply of the symbiont.

### Deciphering the role of nutrient cycling in cnidarian holobionts

Our results suggest (I.) that nutrient cycling is drastically altered between symbiotic and non-symbiotic Aiptasia; (II.) that different *Symbiodinium* types possess different metabolic capabilities within the same Aiptasia strain and (III.) that different Aiptasia strains affect the metabolic performance of the same algal symbiont. Although our results require further validation with regard to their wider applicability beyond the Aiptasia model system, our findings showcase the distinct advantages of a model system approach for the study of nutrient cycling in the the cnidarian – dinoflagellate symbiosis. However, questions remain regarding the precise nature of nutrients exchanged in this symbiosis and the underlying processes involved. Since nutrient exchange is arguably the functional basis of mutualistic association (7), providing answers to these questions is one of the keys to understanding holobiont functioning (14). Furthermore, nutrient cycling is likely a dominant driver of holobiont fitness under varying environmental conditions (58–61). Understanding environmental controls of nutrient cycling may therefore help to provide novel insights on the mechanisms of symbiosis establishment, maintenance, and disruption.

In this context, we formulate three important questions that are relevant for future studies of nutrient cycling in cnidarian holobionts:

I. **How does symbiont diversity affect nutrient exchange within the symbiosis and how does it influence holobiont success under varying environmental conditions?** It has been previously observed that the performance of symbionts depends on the environmental conditions (e.g. during coral bleaching) (62). While many studies have investigated the role of oxidative stress in these phenomena (63, 64), nutrient cycling is likely another important factor involved.
II. **How is nutrient cycling regulated during symbiosis establishment and maintenance?** In contrast to mature coral holobionts, carbon translocation by symbionts appears to be negligible in early stages of symbiosis establishment (51, 65). Understanding the processes around initiating and stabilizing this nutrient exchange during symbiosis development will advance our understanding of the factors underlying the success of this symbiosis.
III. **What is the role of bacteria and other microbes in holobiont nutrient cycling?** It is widely acknowledged that carbon, nitrogen, and sulfur cycling microbes are ubiquitous members of the cnidarian microbiome (32, 66, 67). However, questions remain regarding their relevance and contribution to holobiont function. Studying nutrient exchange between these microbes and other members of the holobiont is necessary to evaluate the importance of the microbiome for holobiont fitness. Future research efforts incorporating a model system approach with field-based coral studies, will transform our understanding of the mechanisms underlying this symbiosis and may prompt new solutions to prevent further loss and degradation of reef ecosystem.

## Author contributions

NR, MA and CRV conceived and designed the experiment. NR, JBR, MP and GP conducted the experiment. PG and MRK carried out NanoSIMS data acquisition. All authors wrote, revised and approved the manuscript.

## Acknowledgements

The authors would like to thank Dr. Rachid Sougrat and Ptissam Bergam from the KAUST imaging core lab for their help with sample preparation. CRV and NR acknowledge funding from the KAUST AIMS CPF partnership funding. Further, research in this publication was supported by KAUST baseline research funds to CRV. The authors would like to acknowledge the Australian Microscopy & Microanalysis Research Facility, AuScope, the Science and Industry Endowment Fund, and the State Government of Western Australian for contributing to the Ion Probe Facility at the Centre for Microscopy, Characterisation and Analysis at the University of Western Australia. JBR was supported by Australian Research Council fellowship DE160100636.

